# Mitochondria-nucleus communication dictates progressive versus bimodal aging of the mammary gland

**DOI:** 10.64898/2026.01.21.700824

**Authors:** Edmund Charles Jenkins, Mrittika Chattopadhyay, Thelma Mashaka, Dmitry Polushakov, Miguel Torres-Martin, Daniela Sia, Igor Bado, Doris Germain

**Author notes:** E.C. J and M.C contributed equally to this study.

## Abstract

Non-linear aging, identified through oscillations in human plasma proteomics around 44 and 60, is postulated to contribute to the onset of diseases. Incidence of breast cancer follows such bimodal pattern. We show that mitochondria-nucleus matching in mice results in two mammary gland aging patterns, including a bimodal pattern associated with susceptibility to mammary tumor and that this bimodal signature is enriched in breast cancer patients diagnosed at 45 and 65.

---

Non-linear aging, identified using human plasma proteomics, is supported by the observation of waves of changes around the ages of 44 and 60^1–3^. However, how these waves are associated with specific diseases remain unclear. Since breast cancer shows a bimodal incidence at 45 and 65 ^4–6^, breast cancer stands out as a disease that may be directly affected by non-linear aging.

Mitochondrial dysfunction is a hallmark of aging ^7–11^. Mitochondria-nucleus matching has profound impact on the resulting nuclear gene expression in several models, including in mice with the C57BL/6 nuclear genetic background (BL/6) but differ in mitochondria genomes, and carry either the C57BL/6 or NZB/OlaHsd (^C57^ or ^NZB^) mitochondria ^12–17^.

We therefore analyzed the aging of the mammary gland using these mice over the entire range of the estrous cycle over aging ^18^.

We aged BL/6^C57^ and BL/6^NZB^ female mice at 3, 6, 11, 14 and 19 months (Extended data 1) based on the reported number of estrous cycles at these ages ^18^ and therefore were deemed to best recapitulate the equivalent of the menopausal status with aging in humans and include pre-menopause, peri-menopause and post-menopause status.

We found that the BL/6^C57^ females, the number of mammary branches (Fig. 1a, b), thickness (Fig. 1d), adipocytes (Fig. 1f, g), collagen (Fig. 1i, j, k) either decreased or showed not significant difference over aging. Tumor formation however, showed a progressive increase with age (Fig. 1o).

**Figure 1:**
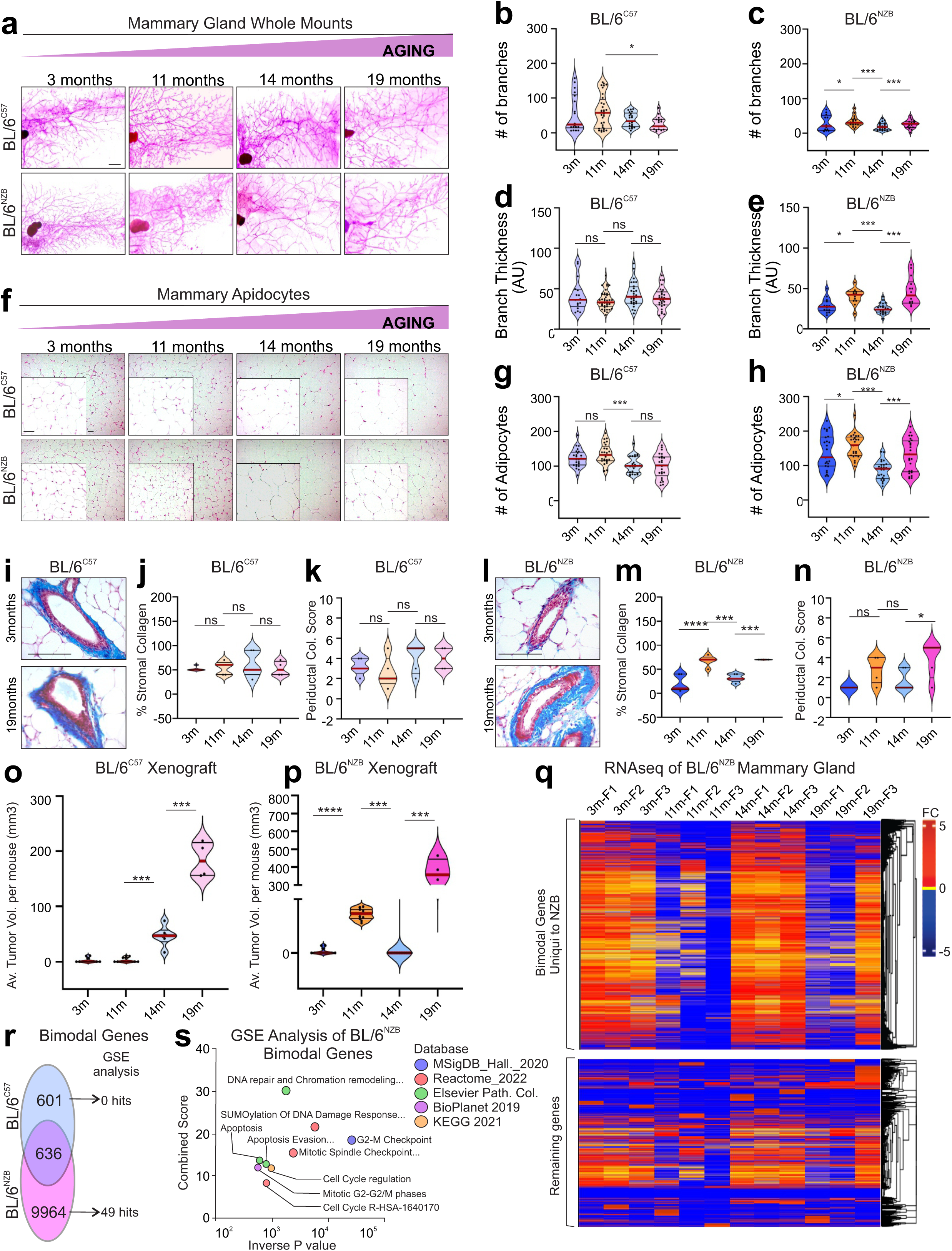
Mammary glands show non-linear aging in BL/6^NZB^ mice. **a**) Mammary gland whole mounts of either BL/6^C57^ or BL/6^NZB^ mice at indicated age. **b-c**) Quantification of the number of mammary branch points (>3). n=8 mice at 3 months and n=5 at other time points and dots indicate 5 area per mouse. **d-e**) Quantification branch thickness (Arbitrary Units). n=5 mice per time point and dots indicate 5 area per mouse. **f**) Staining of mammary adipocytes of either BL/6^C57^ or BL/6^NZB^ mice at indicated age. **g-h**) Quantification of adipocytes per field n=5 mice per time point and dots indicate 5 area per mouse. Masson’s Trichrome Staining (MTS) for collagen of mammary gland sections from BL/6^C57^ (**i**) or BL/6^NZB^ (**l**) mice. Quantification of stromal collagen from BL/6^C57^ (**j**) or BL/6^NZB^ (**m**) mice at indicated age. n=4 biological replicates. Periductal collagen scoring from BL/6^C57^ (**k**) or BL/6^NZB^ (**n**) mice at indicated age. n=4 mice per time points. EO771 Xenograft tumor volume 3 weeks after injection in BL/6^C57^ (**o**) or BL/6^NZB^ (**p**) females at indicated age. n=5 mice per time points. **q**) Heatmap expression analysis of bulk RNAseq from BL/6^NZB^ mice at indicated age. n=3 mice per time points. **r**) Venn diagram of number of genes whose expression follows an oscillation pattern (up/down/up/down) over the aging time course. **s**) Top 9 (out of 52) most significant terms from Gene Set Enrichment (GSE) analysis of bimodal genes unique to BL/6^NZB^ mice. Statistics were performed with an ANNOVA followed by a Tukey post test to detect differences between groups. * p<0.05, **p<0.005, ***p<0.0005

In contrast, in BL/6^NZB^ females, the number of mammary branches (Fig. 1a, c), thickness (Fig. 1e), adipocytes (Fig. 1f, h), collagen (Fig. 1l, m, n) as well as tumor formation (Fig. 1p), showed a bimodal pattern of being increased at 11 and 19 months relative to 3 and 14 months. The bimodal pattern was confirmed by repeating the analysis at both 6 months and 24 months (Extended data 2). This differential pattern of aging and tumor susceptibility between genotypes did not correlate with fluctuation in estrogen levels or estrous cycle (Extended data 3a, b, c) ruling out the possibility that difference in estrogen levels is the driver of the observed differences in aging patterns of the mammary glands.

Bulk RNAseq analysis revealed a large number of genes showing an oscillation pattern over aging in the BL/6^NZB^ females, while this was observed to a far lesser extent in BL/6^C57^ females (Fig. 1q, Extended data 4). We estimated that 9964 genes adopt a bimodal pattern and are unique to BL/6^NZB^ females, while only 601 bimodal genes were found in BL/6^C57^ females (Fig. 1r). Gene Set Enrichment (GSE) analysis for terms related to tumor suppression and cell cycle regulation of bimodal genes unique to BL/6^NZB^ (Fig. 1s).

The bimodal pattern was also observed using extracellular matrix (ECM) proteomics (fig. 2). In the BL/6^C57^ female mice, principal Component Analysis (PCA) (Fig. 2b) and composition of ECM proteins (Fig. 2c) showed no oscillation and remained relatively stable across ages (Fig. 2g, Extended data 5).

**Figure 2:**
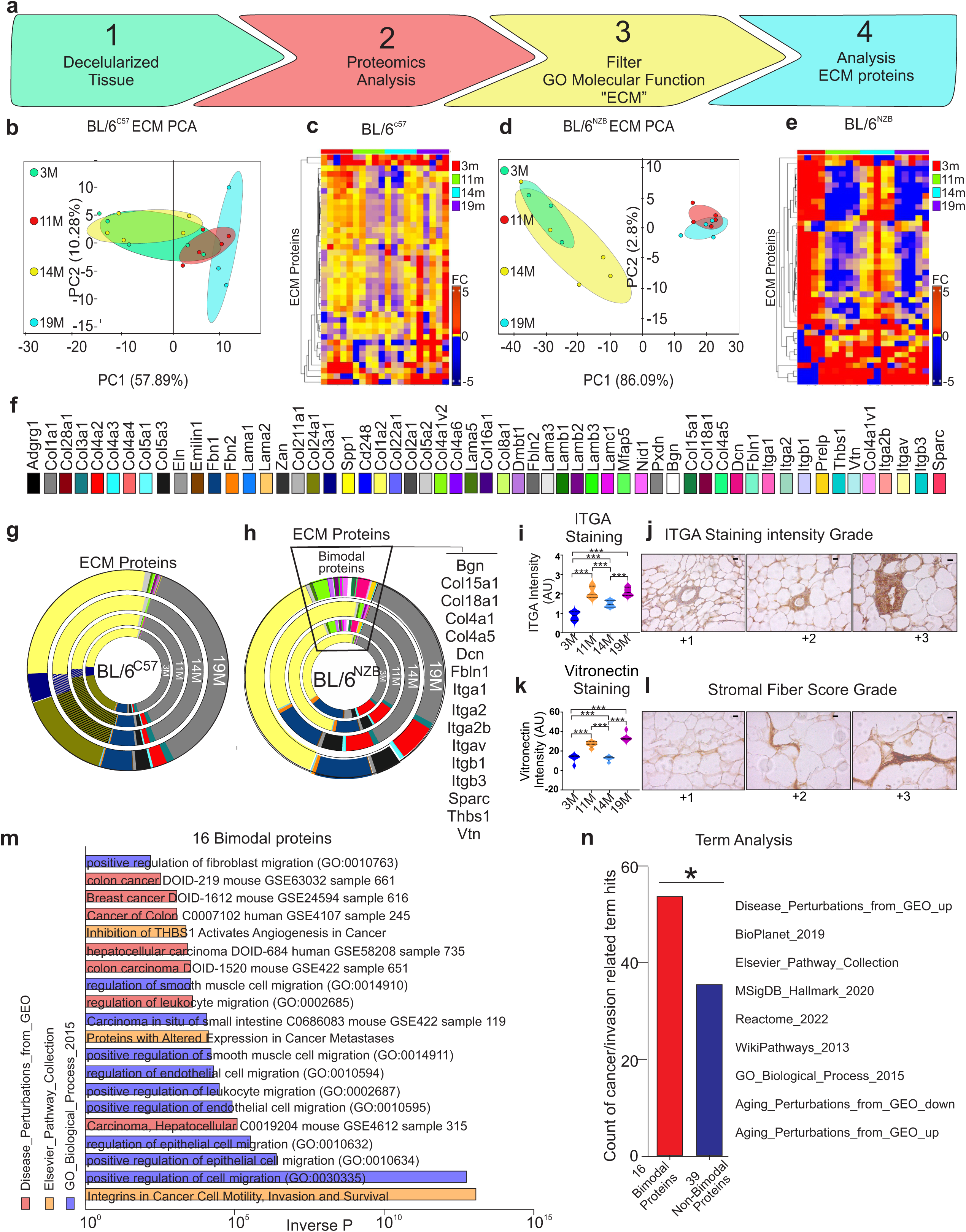
Proteomic analysis of decellularized mammary glands reveals bimodal ECM proteins that are associated with tumor progression in BL/6^NZB^ mice. **a**) Schematic representation of experimental design. Principal component analysis of proteomics analysis from BL/6^C57^ (**b**) or BL/6^NZB^ (**d**) mice at indicated age. n=5 mice per time points. Heatmap of proportional representation of ECM proteins from individual BL/6^C57^ (**c**) and BL/6^NZB^ (**e**) females at indicated ages. **f**) Color code list of ECM proteins used for analysis. Proportional pie graph representation of ECM proteins from BL/6^C57^ (**g**) or BL/6^NZB^ (**h**) mice at 3 indicated age. **i**) Quantification of ITGA staining intensity in BL/6^NZB^ mice at indicated age. n=5 mice per time points and a minimum of 20 ducts/section were used for scoring**. j**) Representative images of ITGA staining and scoring criteria. **k**) Quantification of Vitronectin staining intensity in BL/6^NZB^ mice at indicated age. n=4 mice per time points and a minimum of 20 ducts were used for scoring **l**) representative images of Vitronectin staining and scoring criteria. **m**) Gene Set Enrichment (GSE) analysis of the 16 bimodal ECM proteins indicated in panel e. **n**) Term analysis related to cancer or invasion following GSE analysis of either the 16 bimodalor remaining 39 non-bimodal ECM proteins. *p<0.05 Fisher’s exact t-test.

In contrast, in BL/6^NZB^ females, PCA (Fig. 2d) and heat map (Fig. 2e) analyses revealed clustering between 3 and 14 months, as well as 11 and 19 months of age and identified 16 ECM proteins with a 4-fold or more bimodal change (Fig. 2h, extended data 6). We validated two bimodal ECM proteins by IHC (Fig. 2i, j, k, l) and found that the top 20 pathways uniquely associated with the 16 bimodal proteins relate to cancer (Fig. 2m, n).

We performed single nuclei RNAseq analysis on primary epithelial cells derived from BL/6^NZB^ females (Fig. 3a-f, Extended data 7) and confirmed the bimodal pattern of aging using analysis of cell-type specific transcriptome (Fig. 3g-j, extended data 8). We then search for genes that adopt the bimodal pattern of expression in each cell type and found 79 bimodal genes in the luminal HS cells (extended data 9), 65 bimodal genes in the AV cells (extended data 10), 43 bimodal genes in the ME cells (extended data 11) and 70 bimodal genes in the mixed lineage of HS-AV cells (extended data 12). Pathway analysis of the 79 bimodal genes specifically expressed in the luminal HS cells across multiple databases revealed they are enriched in breast cancer-related pathways (6) and ECM interaction related pathways (10) (Fig. 3k, extended data 13a), while non-bimodal genes in the HS cells showed no ECM related pathways and only one breast cancer related pathway (extended data 13b). Further, analysis of the HS bimodal gene signature in the bulk RNAseq data revealed an oscillation of expression in the BL/6^NZB^ (extended data 14) but not in the BL/6^C57^ females (extended data 15). Gene set analyses of the other cell specific bimodal signatures also reveal cancer related pathways (extended data 16, 17 and 18).

**Figure 3:**
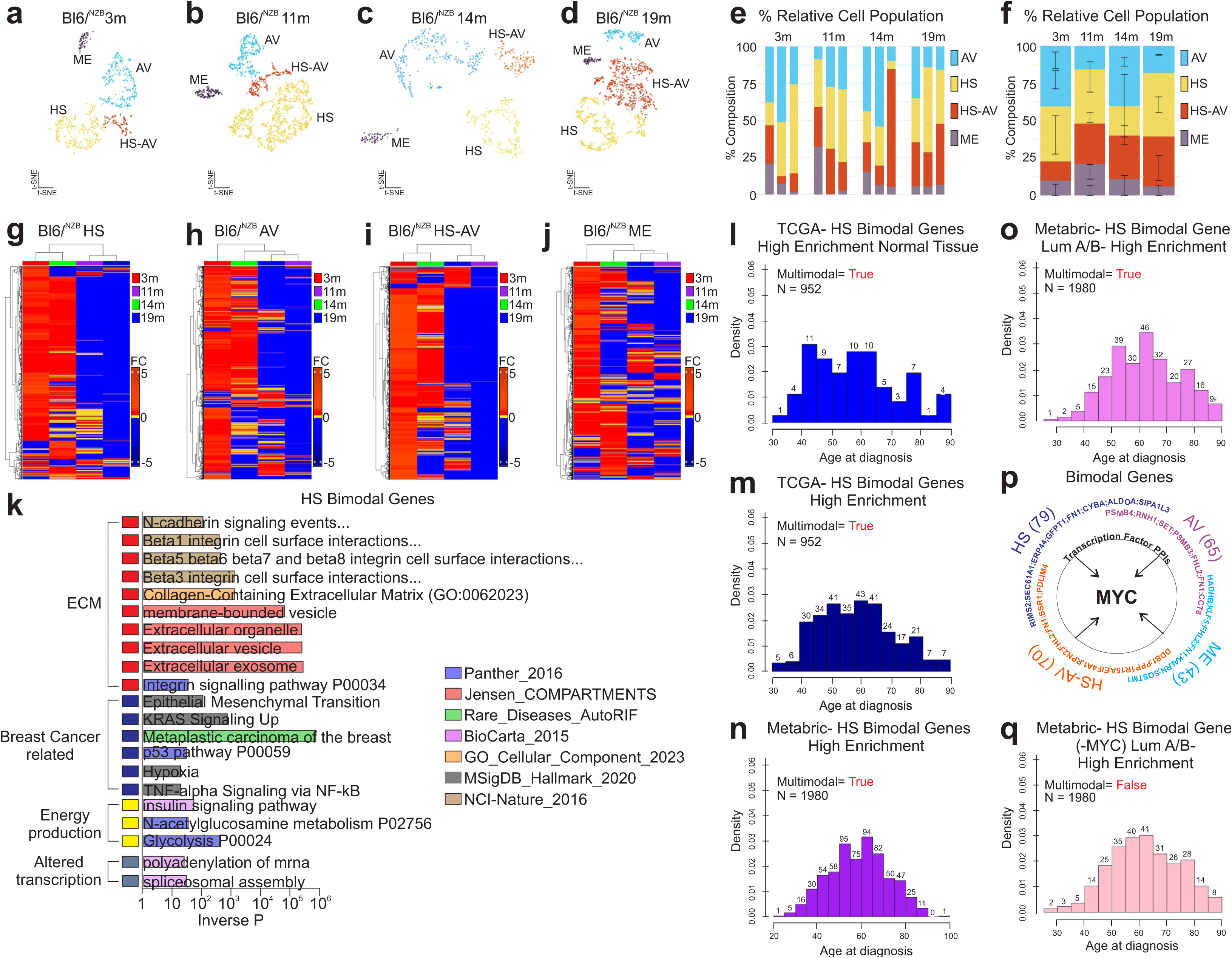
Single nuclei RNAseq analysis shows bimodal fluctuation in the number of individual mammary epithelial cell populations and their transcriptomes in BL/6^NZB^ females over aging. **a**) t-SNE clustergram of single nuclei RNAseq data from 3 BL/6^NZB^ mice at 3 (**a**), 11 (**b**) 14 (**c**), or 19(**d**). HS= Hormone sensitive, HS-AV= hormone sensitive alveolar cells, AV= alveolar cells, ME= myoepithelial cells. **e-f**) Proportional representation of HS, AV, HS-AV, and ME cells in individual mice (e) (n=3 mice per time point, 12 mice total) or combined by time point (f). **g-j**) Heatmap and hierarchical clustering of differential gene expression data at indicated ages in HS cells (g), AV cells (h), HS-AV cells (i), or HS-AV (j). **k**) Gene Set Enrichment analysis of the 79 bimodal genes identified in the HS cell population. **l**) Histogram of patient age at diagnosis for TCGA patients highly enriched for the bimodal gene signature found in HS cells in normal adjacent tissue. **m**) Histogram of patient age at diagnosis for TCGA breast cancer patients highly enriched for the HS bimodal gene signature. **n**) Histogram of patient age at diagnosis for all METABRIC breast cancer patients highly enriched for the HS bimodal gene signature. **o**) Histogram of patient age at diagnosis for Luminal A/B METABRIC breast cancer patients highly enriched for the HS bimodal gene signature. **p**) Protein-Protein Interaction (PPI) analysis of bimodal genes found in HS, AV, HS-AV, and ME cell populations reveals MYC as a common binding partner. **q**) Histogram of patient age at diagnosis for Luminal A/B METABRIC patients highly enriched for the HS bimodal gene signature minus MYC interacting proteins. Multimodality was assessed using a custom density-based test, where a kernel density estimate (bandwidth = ’SJ’) identified significant peaks (local maxima) exceeding a height threshold of 75% of the maximum density, indicating multimodality if multiple peaks were detected. Density values are shown, with counts labeled above each bin.

Since the incidence of breast cancer shows a bimodal age distribution at 45 and 65 ^4–6^ (the equivalent of 11 months and 19 months in mice), if applicable to human, the bimodal gene signatures is predicted to be enriched in patients diagnosed around the age of 45 and 65 years and to be found in the normal adjacent breast tissue since the signature in mice was identified using normal mammary glands. We identified 108 patients in the TCGA dataset for which data is available for both the tumor and normal adjacent tissue from the same patient. We performed single-sample gene set enrichment analysis (ssGSEA) to generate an enrichment score for each patient in the TCGA-BRCA cohort (primary tumor samples) for the HS bimodal gene signature, identified in mice. We divided the cohort into High enrichment score (top tertile) and Low enrichment score (bottom tertile) groups.

Analysis of patient age at diagnosis among normal adjacent tissue samples (Fig. 3l) and in highly enriched patients (Fig.3m) revealed a multimodal distribution with peaks at 45 and 65, while the age distribution of patients classified within Low enrichment score show a unimodal distribution with a peak at 65 (Extended data 19). Patients with High enrichment score were more likely to be younger but not associated with BRCA1 mutation (< or equal to 43 y.o.) (Extended data 20).

To further validate this result, we repeated the analysis in the METABRIC breast cancer dataset and found that in this larger dataset the same multimodal distribution is observed (Fig. 3n, o).

Finally, to gain insight into what may be driving these oscillations, we performed enrichment analysis using the Transcription_Factor_PPIs library in Enrichr and identified the transcription factor MYC (Fig. 3p). Removal of the 13 c-Myc related genes from the HS bimodal genes signature abolished the multimodal distribution (Fig. 3q, Extended data 9).

Collectively, our data suggest that the way breasts age may impact their susceptibility to develop breast cancer and contribute to the bimodal distribution of this disease. However, since two-thirds of breast cancers occur in women over the age of 50 ^19^, we propose that the increased incidence of tumor formation at 65, relative to 45, may be due to the additive effect of bimodal and progressive aging patterns.

## Materials and Methods

### Mice

All mouse experiments were conducted per an approved protocol by the Institutional Animal Care and Use Committee (IACUC). BL/6^C57^ and BL/6^NZB^ mice were originally generated and kindly provided by Dr. José Antonio Enríquez in the C57BL/6JOlaHsd background. The mice were housed in vivariums at the Icahn School of Medicine at Mount Sinai with *ad libitum* access to food and water. BL/6^C57^ and BL/6^NZB^ female mice were aged and harvested at specified time points (n=5 per timepoint for each genotype), and mammary glands were harvested and stored appropriately for subsequent analysis. The estrous cycle stage of all female mice was determined prior to experimentation. To collect vaginal epithelial cells, approximately 50 µl of phosphate-buffered saline (PBS) was aspirated into a pipette tip and gently inserted into the vaginal opening. The PBS was flushed in and out of the vaginal canal two to three times to collect cells. The resulting cell suspension was then evenly distributed onto three microscope slides, forming a thin smear, and left to air dry. Slides were subsequently stained using Modified Giemsa Stain (Sigma-Aldrich, GS500-500ml) for 2–3 minutes, followed by a rinse with water. The slides were examined under a microscope to determine the estrous cycle stage as described^20,21^.

The E0771 murine mammary carcinoma cell line, originally derived from a spontaneous mammary tumor in C57BL/6 mice, was used to establish orthotopic tumors in the mammary glands of mice, following a previously published protocol with minor modifications ^21^. A total of 1 × 10⁵ E0771 cells, suspended in a 1:1 mixture of Hank’s Balanced Salt Solution (HBSS) and growth factor-reduced Matrigel (GIBCO) in a final volume of 50 µl, were injected into the bilateral inguinal mammary glands of estrus-stage-matched (Estrus phase) virgin female mice from two strains: BL/6^C57^ and BL/6^NZB^. Injections were performed at 3, 6, 11, and 19 months of age. Tumor growth was monitored three times per week using electronic calipers, measuring the longest (length) and widest (width) tumor diameters. Tumor volume was calculated using the modified ellipsoid formula: volume = ½ × (length × width²) as described previously ^21^

For measurement of Estradiol levels in mice serum, blood was collected from estrus cycle matched (Estrus phase) virgin mice at different ages. Blood from pregnant mice (late stage) was used a positive control. Serum was separated from all samples and estradiol levels were measured using the Elisa Kit from Enzo (ADI-900-174).

### Wholemounts

Mouse abdominal mammary glands were harvested, mounted on microscope slides, and air-dried briefly. The glands were then fixed in Carnoy’s fixative (75% Ethanol, 25% glacial acetic acid) overnight. Subsequently, they were washed in 70% ethanol for 15 minutes, followed by a 5-minute water rinse, and stained overnight in carmine alum solution (1g carmine (Sigma C1022) and 2.5g aluminum potassium sulfate (Sigma A7167) in 500ml dH_2_O) according to standard protocol. After staining, the carmine-stained whole mounts were rinsed in MilliQ-filtered water for 5 minutes and dehydrated through an alcohol series (70%, 95%,100% EtOH, 15 minutes for each step). The glands were cleared in xylene and mounted using a Permount mounting medium. Finally, the whole mounts were imaged using a Zeiss Stemi 300 microscope.

### Mammary branching assessment

To quantify branching in mouse mammary gland whole-mount images, we developed an image processing pipeline using ImageJ/FIJI (v1.54f). GRB images were opened and split into individual channels to isolate the channel containing the highest contrast between duct and surrounding tissue. Background subtraction was performed by subtracting one channel from another, followed by normalization through division by the red channel twice, both in 32-bit format, to correct for uneven staining. Shading correction was applied using a second-degree polynomial fit in both x and y directions with a regularization parameter of 2 to address illumination artifacts. The resulting image was converted to 8-bit, and contrast was enhanced with a saturation limit of 0.35%.

For segmentation, the Yen thresholding method was applied to generate a binary mask, with the background set to black. Morphological operations were used to refine the mask: dilation (twice) to fill gaps, de-speckling (four times) to remove noise, and erosion (twice) to restore structure boundaries. The binary image was skeletonized to reduce mammary gland structures to single-pixel-wide centerlines. Branching morphology was quantified using the "Analyze Skeleton (2D/3D)" plugin, with no pruning, to measure branch number, branch lengths, junction points, and endpoints. Results were saved as TIFF images and quantitative data tables. The pipeline was automated using an ImageJ macro to ensure reproducibility across samples.

### H&E, Masson Trichome Stain and Immunohistochemistry

Mammary glands were fixed in 10% formalin and subsequently processed and paraffin-embedded for sectioning by the Mt Sinai Biorepository Core Facility. Formalin Fixed Paraffin Embedded (FFPE) sections were prepared and stained for hematoxylin and eosin (H&E) according to standard protocol. Additional unstained slides were processed for collagen using the Fisher Scientific Epredia Richard-Allan Scientific Masson Trichrome Kit following the manufacturer’s instructions. Stained slides were imaged using a Zeiss AX10 Microscope. Quantification of collagen content was performed by calculating the percentage of stromal collagen per mammary gland and assessing periductal thickness using an arbitrary scoring system ranging from 1(thin/filamentous collagen) to 5 (thick collagen). This scoring was conducted by a blinded observer.

For immunohistochemical analysis, tissue sections were first deparaffinized by immersing them in xylene twice for 5 minutes each. This was followed by gradual rehydration through a descending ethanol gradient (100%, 90%, and 70%), ending with distilled water. Antigen retrieval was performed by incubating the sections in 10 mM sodium citrate buffer (pH 6.0) at 90–100°C for 30 minutes. Slides were then allowed to cool gradually to room temperature and rinsed twice with Tris-buffered saline (TBS). Immunostaining was carried out following the manufacturer’s protocol for the ImmPRESS Excel Amplified Polymer Kit (Vector Laboratories, Cat# MP-7601 and MP-7602). Primary antibodies against Vitronectin (Abcam, Cat# AB235987) and ITGA11 (Abcam, Cat# AB316249) were applied at a 1:100 dilution and incubated overnight at room temperature in a humidified chamber.

### Mammary gland decellularization

Freshly dissected mammary glands were decellularized by immersion in 1% SDS in TBS containing penicillin/streptomycin and DNase (1U/mL) in a 15 mL Falcon tube. The tubes were shaken at room temperature for 48 hours with 6 complete changes of the decellularization solution. Subsequently, the glands underwent an additional 48-hour period with 6 complete changes of water, (as described by Jenkins et.al., 2022). After decellularization, the mammary glands were frozen in 10% DMSO in TBS to preserve the tissue integrity before the samples were sent for proteomics analysis.

### Proteomics

#### Protein digestion

Decellularized mammary glands were resuspended in 50 uL of 8M Urea, 50mM EPPS and treated with DL-dithiothreitol (5mM final concentration) for 2 hours at 37°C with shaking (1400 rpm) on a Thermomixer (Thermo Fisher). Free cysteine residues were alkylated with 2-iodoacetamide (10mM final concentration) for 30 minutes at 25°C in the dark. Urea was diluted to 2M with Ammonium Bicarbonate (ABC) 100mM and PGNase added for 2 hours 37°C with shaking (1400 rpm). LysC (1:100 enzyme:protein) was added, followed by incubation for 2h at 37°C at 1400 rpm. Finally, trypsin (1:100 enzyme:protein) was added, followed by overnight incubation at 37°C at 1400 rpm. After the overnight incubation, the digest was acidified to pH <3 with the addition of 50% trifluoroacetic acid (TFA), and the peptides were desalted on 3-plug C18 (3M Empore^TM^ high performance extraction disks) stage tips. Briefly, the stage tips were conditioned by sequential addition of i) 100 μL Methanol, ii) 100 μL 70% Acetonitrile (ACN)/0.1% TFA, iii) 100 μL 0.1% TFA twice. Following conditioning, the acidified peptide digest was loaded onto the stage tip. The stationary phase was washed once with 100 μL of 0.1% Formic Acid (FA). Finally, samples were eluted using 50μL of 70% ACN/0.1% FA twice. Eluted peptides were dried under vacuum followed by reconstitution in 12 μL of 0.1% FA, sonication and transfer to an autosampler vial. Peptide yield was quantified by NanoDrop (Biotek Synergy H1).

### MS analyses

Peptides were separated on a 50 cm column composed of C18 stationary phase (Thermo Fisher ES903) using a gradient from 0.5% to 25% buffer B over 100 minutes, to 50% over 15 minutes, to 90% in 5 minutes (buffer A: 0.1% FA in HPLC grade water; buffer B: 99.9% ACN, 0.1% FA) with a flow rate of 300nL/min using a nanoAcquity Hplc system (Waters). MS data were acquired on a Eclipse mass spectrometer (Thermo Fisher Scientific) using a data-independent acquisition (DIA) method. The method consisted of one MS1 scan, 400000 AGC target, maximum injection time of 50 msec, scan range of 380-985 m/z and a resolution of 120K. Fragment ions were analyzed in 60 DIA windows, maximum injection time of 40 msec, 100000 AGC target at a resolution of 15K.

### DIA Data Analysis

Raw data files were processed using Spectronaut version 18.5 (Biognosys) and searched with the PULSAR search engine with a mouse UniProt protein database downloaded on 2022/09/26 (94,765 entries). cysteine carbamidomethylation was specified as fixed modifications, while Methionine oxidation, acetylation of the protein N-terminus and Deamidation (NQ) were set as variable modification. A maximum of two trypsin missed cleavages were permitted. Searches used a reversed sequence decoy strategy to control peptide false discovery rate (FDR) and 1% FDR was set as threshold for identification. Unpaired t-test was used to calculate p-value in differential analysis, volcano plot was generated based on log2FC and q-value (multiple testing corrected p-value using Benjamini-Hochberg method). A q-value of ≤0.05 was considered the statistically significant cut-off.

### Bulk RNA sequencing

Mammary glands of BL/6^C57^ and BL/6^NZB^ females at 3, 11, 14 and 16 months of age were flash frozen on liquid nitrogen and sent to Azenta for bulk RNA sequencing. The RNA sequencing data was analyzed using BioJupies, an interactive online platform that automates the generation of principal component analysis, hierarchical clustering, and differential gene expression analysis. Gene Set Enrichment Analysis was performed using Enrichr, focusing on the top differentially expressed genes and identifying enriched biological pathways associated with the dataset.

### Single Cell RNA Sequencing and Analysis

Freshly isolated snap frozen Inguinal mammary glands were submitted to GENEWIZ, LLC, NJ, USA for nuclei isolation and GEM (Gel Bead-based Emulsion) single cell RNA seq analysis on the 10x Genomics platform. Cellranger 7.0.1 was used to perform alignment of the raw FASTQ reads, filtering, barcode counting, UMI counting, and generation of the cloupe files used in downstream analysis. The cellranger count command was run with default settings, aligning to the mm10-2020-A Mus musculus 10x reference genome. The loupe browser was used to identify cells expressing markers published by Brugge and are as follow: Mfge8, Trf, Csn3, Wfdc18, Ltf (Luminal Alveolar (AV) cells), Prlr, Pgr, Esr1, Cited1, Prom1 (Luminal Hormone Sensitive (HS) cells), Krt17, Krt14, and Krt5 (Myoepithelial cells). This browser feature is described here (https://support.10xgenomics.com/single-cell-atac/software/visualization/latest/tutorial-celltypes). No additional code was used. Azenta Life Sciences uses the 10x Genomics Chromium Single Cell Gene Expression kits, which include options for nuclei isolation. Specifically, they have optimized workflows for processing both cells and nuclei with the 10x Genomics Chromium platform. 10x Genomics Chromium Single Cell Gene Expression Flex Kit. This kit is designed for flexibility, allowing the processing of both cells and nuclei. Suitable for a broader range of sample types, including fresh or frozen cells, nuclei from various tissues, and even samples with low viability. The Flex Kit supports the processing of both single-cell and single-nuclei RNA sequencing in one workflow but requires additional steps or reagents for nuclei isolation if that’s the primary focus. After using the 10x Genomics Chromium Single Cell Gene Expression Flex Kit for nuclei or cell isolation, Azenta Life Sciences follows a detailed library preparation process. After isolating nuclei using the Flex Kit or the Nuclei were reuspended in a suitable buffer.The were combined with barcoded Gel Beads, Master Mix, and Partitioning Oil in the 10x Chromium Controller or Next GEM Chip. This step encapsulates individual nuclei or cells in Gel Beads-in-emulsion (GEMs), where each GEM contains one nucleus or cell along with a unique barcode.Reverse transcription occurs inside the GEMs, converting the RNA into barcoded cDNA. cDNA is amplified through PCR. The amplified cDNA was fragmented to an appropriate size for sequencing. The ends of the fragmented cDNA were repaired, and an ’A’ base is added to facilitate adapter ligation. Unique Dual Index (UDI) Adapters: Adapters with sample-specific indices were ligated to the cDNA fragments. The library with the sample index adapters was then amplified. A Bioanalyzer was used to check the size distribution and quality of the library, ensuring only high-quality libraries proceed to sequencing.

### Statistics

Statistical analyses were conducted using GraphPad Prism Software. A p-value cutoff of 0.05 was applied to determine significance levels. ns = not significant, * p < 0.05, ** p < 0.01, *** p < 0.001, **** p < 0.0001.

Correlations of high and low signature patient groups with clinico-pathological characteristics were analyzed by Fisher’s exact test.

## Acknowledgments

We thank Dr. Mara Monetti, director of the Proteomic Core facility to Memorial Sloan Kettering to perform the mass spectrometry analysis of ECM proteomics. This work is supported by an NIH R01AG075589 to D.G and a grant from the Samuel Waxman Cancer Research Foundation.

## Contributions

E.C.J analyzed the proteomics, bulk and single cell RNAseq data and has generated data in figures 1q, r, s, 2c, e, g, h, m, n and all of figure 3. He also generated all supplementary figures 4, 5, 6, 7, 8, 9, 10, 11, 12, 13, 14, 15, 16, 17, 18, 19. M.C has generated data included in figure 1a-p, figure 2i-l, supplementary figs. 1, 2 and 3 and helped colony management and mammary gland analysis. T.M has generated data included in figure 1a-p and supplementary figure 1a,b, c. D.P has helped in the analysis of the proteomics data in figure 2f.

M. M is a bioinformatician that contributed to the analysis in figure 3. D.S has generated the enrichment score and perform some of the analysis in human data in figure 3 as well as supplementary figure 20. I.B has provided access the METABRIC dataset. D.G has guided the project, interpreted the data and wrote the manuscript.

## Peer review

### Peer review information

*Nature Aging* thanks the anonymous reviewers for their contribution to the peer review of this work.

## Additional information

**Publisher’s note** Springer Nature remains neutral with regard to jurisdictional claims in published maps and institutional affiliations.

## Extended data

**Extended data 1:** Experimental design and analysis of mammary gland at base line at 3 and 6 months.

**Extended data 2:** Tumor formation at 6 and 24 months.

**Extended data 3:** Estrogen levels an estrous cycle.

**Extended data 4:** Heatmap of bulk RNAseq in BL/6^C57^ females.

**Extended data 5:** Average proportional values of ECM proteins measured by proteomics analysis from BL6/^C57^ mice at 3,11, 14, or 19 months of age

**Extended data 6:** Average proportional values of ECM proteins measured by proteomics analysis from BL6/^NZB^ mice at 3,11, 14, or 19 months of age

**Extended data 7:** Violin plots of the Log2 Sum Expression values of the markers used for snRNAseq analysis.

**Extended data 8:** Heatmaps and hierarchical clustering of snRNAseq of individual mice for each cell population.

**Extended data 9:** List of bimodal genes in HS cell population.

**Extended data 10:** List of bimodal genes in AV cell population.

**Extended data 11:** List of bimodal genes in ME cell population.

**Extended data 12:** List of bimodal genes in HS-AV cell population.

**Extended data 13:** GSE analysis of bimodal and non-bimodal genes in HS cells.

**Extended data 14:** Heatmap of the HS bimodal genes signature in the bulk RNAseq data from BL/6^NZB^ mice.

**Extended data 15:** Heatmap of the HS bimodal genes signature in the bulk RNAseq data from BL/6^C57^ mice.

**Extended data 16:** GSE analysis of bimodal and non-bimodal genes in AV cells.

**Extended data 17:** GSE analysis of bimodal and non-bimodal genes in HS-AV cells.

**Extended data 18:** GSE analysis of bimodal and non-bimodal genes in ME cells.

**Extended data 19:** Histogram of distribution of patients with bottom tertile HS signature

**Extended data 20.** Table of comparison of clinical characteristics between patients with high and low enrichment score of the 79 genes signature.

## Source data

The single cell RNAseq data is available (GSE275925) and the raw files of bulk RNSseq can be found at the SRA project PRJNA1153611.

## Rights and permissions

Springer Nature or its licensor (e.g. a society or other partner) holds exclusive rights to this article under a publishing agreement with the author(s) or other rightsholder(s); author self-archiving of the accepted manuscript version of this article is solely governed by the terms of such publishing agreement and applicable law.

